# Quantitative Assessment of shRNA Loading and Delivery Efficiency of Engineered Extracellular Vesicles

**DOI:** 10.64898/2026.06.05.730346

**Authors:** Julia A. Rädler, Giulia Corso, Omnia Elsharkasy, Noriyasu Kamei, Doste R. Mamand, Xiuming Liang, Wenyi Zheng, Antje M. Zickler, Houze Zhou, Samantha Roudi, Oscar P.B. Wiklander, Imre Mäger, Dhanu Gupta, Samir EL Andaloussi

## Abstract

RNA interference (RNAi) therapeutics enable selective silencing of disease-associated genes. Yet, their clinical application remains largely confined to the liver due to extrahepatic delivery constraints of current platforms such as GalNAc conjugates and lipid nanoparticles. Extracellular vesicles (EVs) offer an attractive alternative delivery strategy owing to their biocompatibility, ability to traverse biological barriers, and amenability to engineering. However, EV-mediated RNA delivery is limited by inefficient endogenous RNA loading and poor cytosolic release following uptake. Here, we establish a modular EV-based platform that addresses both challenges by integrating enhanced endogenous shRNA loading with fusogen-mediated cytosolic delivery. Using Argonaute 2 (AGO2)-assisted loading, we substantially increase shRNA copy numbers per vesicle (up to 3.7 copies/EV) and enable quantitative, molecule-resolved assessment of delivery potency. Engineered EVs achieve robust and reproducible shRNA-mediated gene silencing with picomolar IC_50_ values across multiple cell types and induce significant target knockdown in the mouse brain following intracerebral administration. Together, these findings demonstrate that coordinated engineering of shRNA loading and cytosolic release can overcome key limitations of EV-mediated small RNA delivery.

## Introduction

RNA interference (RNAi) enables post-transcriptional gene silencing through short non-coding RNAs and has emerged as a powerful therapeutic modality (1, 2). In mammalian cells, microRNAs (miRNAs) and short-hairpin RNAs (shRNAs) are transcribed in the nucleus and processed through RNAi biogenesis pathways, whereas small-interfering RNAs (siRNAs) typically refers to synthetic double-stranded RNA in therapeutic contexts (3, 4). These RNA species associate with Argonaute (AGO) proteins to form the RNA induced silencing complex (RISC), which represses target mRNAs through cleavage or translational inhibition (5). The ability of RNAi to modulate virtually any gene has driven the development of RNAi-based therapeutics, several of which have now received regulatory approval, with many more progressing through late-stage clinical trials (6). However, the hydrophilic nature and poor bioavailability of naked RNAi agents require the use of efficient delivery systems. Current siRNA delivery platforms rely predominantly on ligand conjugates, such as N-acetylgalactosamine (GalNAc), or encapsulation within lipid nanoparticles (LNPs) (7). While these technologies have demonstrated clinical success, both are characterized by preferential hepatic accumulation due to intrinsic biodistribution and receptor expression patterns, thereby limiting applications beyond liver-associated diseases.

Extracellular vesicles (EVs) have emerged as a promising alternative delivery platform. These nanoscale, membrane-enclosed vesicles are secreted by all cell types and mediate intercellular communication by transferring proteins, lipids, and nucleic acids (8, 9). Their endogenous origin, biocompatibility, ability to cross biological barriers, and amenability to molecular engineering have spurred interest in exploiting EVs for therapeutic cargo delivery (10, 11). EVs have shown favorable safety profiles in early-phase clinical studies and are currently being evaluated for RNA delivery applications, including the delivery of siRNA targeting mutant *KRAS* in pancreatic cancer (12–14).

Despite their promise, EVs face inherent limitations as RNA delivery vehicles. RNA copy numbers per vesicle are typically low (15, 16), and functional delivery is further limited by inefficient cytosolic access, as internalized cargo often remains sequestered within endo-lysosomal compartments following uptake (13). Moreover, most studies investigating EV-mediated delivery of small RNAs have relied on exogenous loading approaches, such as electroporation or chemical transfection into isolated vesicles. However, these methods often result in heterogeneous RNA incorporation, cargo aggregation, and limited reproducibility (17, 18). On the other hand, endogenous loading strategies, which exploit cellular sorting mechanisms during EV biogenesis for biomolecular engineering, offer a promising alternative. Accumulating evidence indicates that RNA incorporation into EVs is selective and regulated, rather than a passive reflection of cytoplasmic abundance (19). For small RNAs, preferential sorting into EVs has been linked to defined sequence motifs, secondary structure, cellular localization, and specific interactions with RNA-binding proteins (RBPs) (20). Although the molecular details of these pathways remain to be fully resolved, these studies collectively suggest that rational manipulation of RNA intrinsic features and RBP engagement may provide a route to enhance endogenous loading of small RNAs into EVs. Importantly, endogenous loading is compatible with complementary EV engineering strategies, such as incorporation of fusogenic membrane proteins during EV biogenesis to promote endosomal escape and enhance functional delivery (21).

In this study, we establish a modular EV-based platform for shRNA delivery that integrates enhanced endogenous RNA loading with fusogen-mediated cytosolic release. Quantitative analysis of RNA copy number and functional potency reveals how RNA loading, cytosolic release, and host RNAi involvement contribute to effective EV-mediated RNA delivery, resulting in robust target knockdown *in vitro* and in the mouse brain *in vivo*.

## Materials and methods

### Construct design and generation

shRNAs were designed using online design tools provided by Biosettia (shRNA Designer) and the Broad Institute Genetic Perturbation Platform (GPP) web portal. All shRNA constructs followed the same structural design, consisting of a KpnI restriction site, the sense strand, a loop sequence (TTCAAGAGA), the antisense strand, a termination signal (TTTTTT), and a NotI restriction site for cloning into a guide RNA expression vector under the control of the U6 promoter.

**Table 1:**
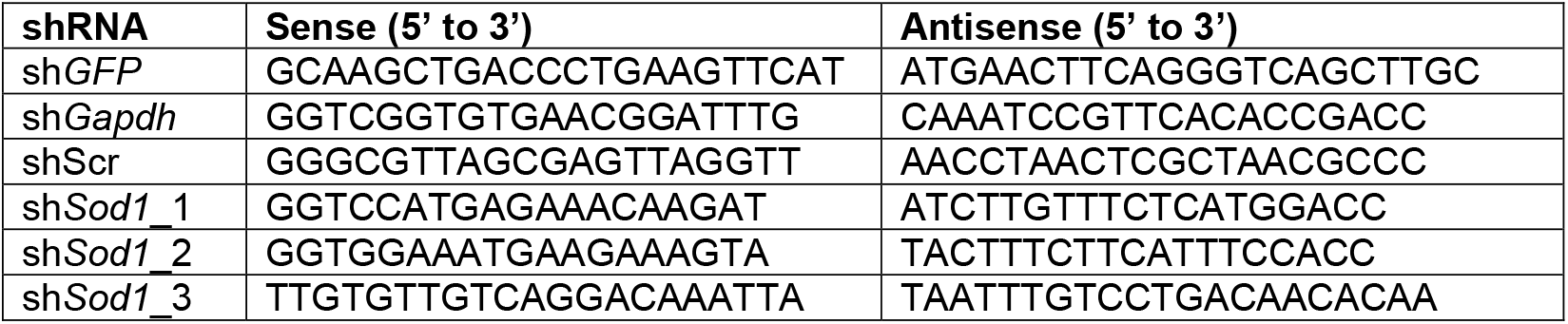
shRNAs used in this study. Nucleotide sequences of sense and antisense strands for each design.

The *Sod1*-targeting siRNA (si*Sod1*) was designed based on a previously published sequence, corresponding to design “VI” (22). Sense and antisense strands were synthesized by Axolabs GmbH.

For Ago2 knockdown experiments, a pre-designed siRNA targeting *Ago2* (si*Ago2*; siRNA ID 152967) was obtained from Thermo Fisher Scientific.

All constructs (AGO2, myrAGO2, CD63-AGO2, TSPAN2-AGO2, VSV-G, and RVG) were cloned into the pLEX vector backbone using standard restriction enzyme-based cloning strategies.

A point mutation in AGO2 (D597A; nucleotide change 1790 A→C) was introduced by primer extension mutagenesis using Phusion™ Green Hot Start II HighFidelity PCR Master Mix (Thermo Fisher Scientific, F566L). Successful mutagenesis was confirmed by Sanger sequencing.

### Cell culture

All cell lines were maintained at 37°C in a humidified incubator with 5% CO_2_ unless otherwise specified. HEK293T, Neuro-2a (N2a), NIH/3T3, C2C12, Hepa1-6, and B16F10 cells were cultured in Dulbecco’s Modified Eagle’s Medium, high glucose (DMEM; Gibco, 31966047) supplemented with 10% fetal bovine serum (FBS; Gibco, A5256701) and 1X antibiotic–antimycotic solution (AA; Gibco, 15240062). Reporter HEK293T cells were maintained in the same medium supplemented with puromycin (2 µg/mL; Sigma-Aldrich, P8833) for selection. Mouse embryonic fibroblasts (MEFs) were cultured in DMEM supplemented with 15% FBS and 1X AA. BV2 cells were cultured in RPMI-1640 medium (Gibco, 72400054) supplemented with 10% FBS and 1X AA.

### Extracellular vesicle production

For EV production, 1 × 10^7^ HEK293T were seeded in 15-cm culture dishes in complete growth medium. At 70-80% confluency cells were transfected with plasmids of interest complexed with polyethylenimine “Max” (PEI MAX® -Transfection Grade Linear Polyethylenimine Hydrochloride (MW 40,000), Polysciences, Cat. #24765; 40 μg DNA: 120 μg PEI) in Opti-MEM Reduced Serum Medium (Gibco, Cat. #31985054). For EV production with siRNA, 20 μg DNA and 50 μg siRNA were transfected using 120 μg PEI. 5 h after transfection, medium was changed to Opti-MEM supplemented with 1X AA. Cells were incubated for 48 h before EV isolation.

### Extracellular vesicle isolation and purification

Conditioned medium (CM) from transfected HEK293T cells was collected and sequentially centrifuged at 700 × g for 5 min and 2,000 × g for 10 min to remove cells and cellular debris. The resulting supernatant was filtered through a 0.22 µm polyethersulfone (PES) vacuum filter (Filtermax “rapid”, TPP).

For CM volumes below 100 mL, samples were directly concentrated by ultrafiltration using Amicon® Ultra centrifugal filters with a 100 kDa molecular weight cutoff (MWCO; Millipore, UFC9100) by centrifugation at 4,000 × g and 4°C to a final volume of ∼500 µL. For CM volumes exceeding 100 mL, samples were first concentrated by tangential flow filtration (TFF; MicroKross, Spectrum Labs; 300 kDa MWCO) to a final volume of approximately 30 mL before ultrafiltration as described above.

In both cases, the concentrate was then loaded onto qEVoriginal/70 nm Gen 2 size exclusion chromatography columns (Izon Science, ICO70) equilibrated with PBS. After discarding the initial 3 mL eluate, vesicle containing fractions (3 mL) were collected according to the manufacturer’s instructions. EV fractions were subsequently concentrated by ultrafiltration using Amicon^®^ Ultra centrifugal filters with a 10 kDa MWCO (Millipore, UFC201024) under the same centrifugation conditions to a final volume of ≤100 µL. EV preparations were used immediately or stored at −80°C until further analysis.

### Nanoparticle tracking analysis

Particle size distribution and concentration of isolated EVs were determined by nanoparticle tracking analysis (NTA) using a ZetaView® x30 TWIN instrument (Particle Metrix). EV samples were diluted in 0.22 µm filtered PBS to achieve particle concentrations within the optimal detection range and measured at 11 positions per sample. Measurements were acquired using standard settings, with camera sensitivity set to 80 and shutter speed set to 100. Data were analyzed using the ZetaView software.

### Extracellular vesicle uptake (*in vitro*)

For EV uptake studies assessing target knockdown by flow cytometry, HEK293T GFP reporter cells were seeded in 96-well plates at a density of 1 × 10^4^ cells per well in complete DMEM without puromycin. The following day, EVs were added at the indicated doses (ranging from 1 × 10^7^–1 × 10^10^ particles per well), and cells were incubated for an additional 48 h. GFP expression was subsequently quantified by flow cytometry to assess target knockdown.

For EV uptake studies followed by RNA extraction, cells were seeded in 96-well plates at defined densities. N2a, NIH/3T3, BV2, and Hepa1-6 cells were seeded at 7.5 × 10^3^ cells per well, whereas B16F10, C2C12, and MEF cells were seeded at 5 × 10^3^ cells per well. The following day, EVs were added at the indicated doses (ranging from 1 × 10^7^–1 × 10^10^ particles per well), and cells were incubated for an additional 48 h prior to RNA extraction.

For EV uptake studies followed by protein analysis by Western blot, N2a cells were seeded in 24-well plates at a density of 5 × 10^4^ cells per well. The following day, EVs were added at doses ranging from 5 × 10^7^ to 5 × 10^9^ particles per well. Cells were incubated with EVs for 48 h prior to protein extraction.

### Transfection (*in vitro*)

For plasmid DNA (pDNA) transfection, cells were seeded in either 96-well or 24-well plates at the densities described above and allowed to adhere for 24 h prior to transfection. pDNA transfections were performed using Lipofectamine™ 2000 (Thermo Fisher Scientific) according to the manufacturer’s protocol. In 96-well plates, 100 ng pDNA was transfected per well using 0.2 µL Lipofectamine™ 2000, whereas in 24-well plates, 500 ng pDNA was transfected per well using 1.1 µL Lipofectamine™ 2000. Following transfection, cells were incubated for an additional 48 h prior to downstream analyses.

For *Sod1* siRNA transfection, cells were transfected at a final amount of 1 pmol siRNA per well using Lipofectamine™ RNAiMAX (0.3 µL per well; Thermo Fisher Scientific, 13778150) according to the manufacturer’s instructions. Cells were incubated for 48 h following transfection prior to RNA isolation.

### EV uptake in Ago2-depleted cells

For depletion of Ago2, N2a cells were reverse-transfected with *Ago2* siRNA. Briefly, transfection complexes consisting of 1 pmol *Ago2* siRNA and 0.3 µL Lipofectamine™ RNAiMAX per well were prepared in 96-well plates according to the manufacturer’s instructions prior to cell seeding. N2a cells were then added at a density of 2 × 10^4^ cells per well and incubated for 24 h.

For EV uptake studies in Ago2-depleted cells, the culture medium was replaced with fresh complete DMEM, and EVs were subsequently added at the indicated doses. Cells were incubated with EVs for an additional 48 h prior to downstream analysis.

### Flow cytometry

Two days after EV addition, cells were detached using 0.05% trypsin–EDTA (Thermo Fisher Scientific), which was neutralized with complete culture medium containing 10% fetal bovine serum. Cells were stained with DAPI (25 ng/mL; Thermo Fisher Scientific) to exclude non-viable cells and analyzed by flow cytometry using a MACSQuant^®^ Analyzer 10 (Miltenyi Biotec). Cells were sequentially gated to exclude debris and doublets, followed by selection of viable cells based on DAPI exclusion. GFP knockdown was quantified as the mean fluorescence intensity (MFI) of the GFP positive population. Flow cytometry data were analyzed using FlowJo™ software version 10 (Becton, Dickinson & Company).

### RNA isolation

For RNA isolation from EV recipient cells, culture medium was completely removed and plates were stored at −80 °C until further processing. Total RNA was extracted using the Maxwell^®^ RSC simplyRNA Cells Kit (Promega, AS1390) on a Maxwell^®^ RSC Instrument (Promega), according to the manufacturer’s instructions. RNA concentration was determined using the Qubit™ RNA High Sensitivity Assay (Invitrogen, Thermo Fisher Scientific, Q32855) on a Qubit™ 3 Fluorometer (Invitrogen, Thermo Fisher Scientific).

For RNA isolation from EVs, EV samples (2 × 10^10^ particles) were adjusted to equal volumes with PBS, keeping the final volume below 75 µL as recommended by the manufacturer. RNA was isolated using the Maxwell^®^ RSC miRNA Plasma Kit (Promega, AS1680) and the corresponding miRNA Plasma protocol on the Maxwell^®^ RSC Instrument, according to the manufacturer’s instructions.

For RNA isolation from tissue samples (striatum), tissue was homogenized using a TissueLyser II (QIAGEN) for 10 min at 30 Hz with a 5 mm stainless steel bead (QIAGEN, 69989) in 200 µL homogenization solution. Total RNA was subsequently isolated using the Maxwell^®^ RSC simplyRNA Tissue Kit (Promega, AS1340) according to the manufacturer’s protocol. RNA concentration was measured using a NanoDrop spectral analyzer.

### Gene expression analysis (RT-qPCR)

For cDNA synthesis, 100–200 ng (cells) or 500 ng (striatum) of total RNA were reverse transcribed using the HighCapacity cDNA Reverse Transcription Kit (Applied Biosystems, 4368813) according to the manufacturer’s instructions. Quantitative PCR (qPCR) was performed on a CFX Opus 96 RealTime PCR System (BioRad) using TaqMan™ Fast Advanced Master Mix (Applied Biosystems, 4444557), with 10 ng cDNA used per reaction.

Custom probe-based assays were used for *Gapdh* (forward: GCC TTC CGT GTT CCT ACC; reverse: CCT CAG TGT AGC CCA AGA TG; probe: /5HEX/CGC CTG GAG/ZEN/AAA CCT GCC AAG TA/3IABkFQ/) and *Hprt* (forward: GCC CTC TGT GTG CTC AAG; reverse: CCC CGT TGA CTG ATC ATT ACA; probe: /56FAM/AGC AGG TCA/ZEN/GCA AAG AAC TTA TAG CCC/3IABkFQ/). TaqMan^®^ Gene Expression Assays were used for *Ago2* (Mm03053414_g1), *Sod1* (Mm01344233_g1) and *Gapdh* (Mm99999915_g1).

Relative gene expression levels were calculated using the ΔΔCt method. Expression of *Gapdh* (custom assay) was normalized to *Hprt. Ago2* expression was normalized to *Hprt. Sod1* expression was normalized to *Gapdh* (TaqMan assay). All expression values were further normalized to the untreated control group.

### Protein extraction and Western blot

For protein analysis of EVs, aliquots corresponding to 5 × 10^9^ EVs were mixed with 4× sample buffer and heated at 70 °C for 10 min prior to SDS–PAGE.

For cellular protein extraction, N2a cells previously incubated with EVs in 24-well plates were detached using 0.05% Trypsin-EDTA (Gibco, 25300054), transferred to microcentrifuge tubes, and pelleted by centrifugation at 900 × g for 5 min. Cell pellets were either stored at −80 °C or immediately lysed in RIPA buffer supplemented with a protease inhibitor cocktail. Clarified lysates were obtained by centrifugation at 13,000 × g for 12 min, and the supernatants were transferred to fresh tubes, mixed with 4× sample buffer, and heated at 70 °C for 10 min prior to electrophoresis.

Protein samples were resolved on NuPAGE^®^ Novex® 4–12% BisTris gels (Invitrogen, Thermo Fisher Scientific) and separated by electrophoresis at 120 V for 2 h in NuPAGE® MES SDS running buffer according to the manufacturer’s protocol. Proteins were transferred to nitrocellulose membranes using iBlot^®^ 2 Transfer Stacks and the iBlot^®^ 2 Gel Transfer Device (Invitrogen, Thermo Fisher Scientific) for 7 min.

Membranes were blocked for 1 h at room temperature with Odyssey® Blocking Buffer (LICOR) under gentle agitation and subsequently incubated with primary antibodies diluted in blocking buffer either overnight at 4 °C or for 1 h at room temperature. Primary antibodies used in this study included anti-β-actin (1:10,000; Sigma Aldrich, A5441), anti-Gapdh (1:10,000; Invitrogen, PA116777), anti-AGO2 (1:500; Abcam, ab32381), anti-VSV-G (1:1,000; Thermo Fisher Scientific, PA130278), anti-CD81 (1:200; Santa Cruz Biotechnology, sc9158), and anti-ALIX (1:1,000; Abcam, ab117600).

Following primary antibody incubation, membranes were washed five times for 5 min each with PBS containing 0.1% Tween-20 (PBS-T) and incubated for 1 h at room temperature with IRDye^®^-conjugated secondary antibodies (LI-COR Biosciences) diluted 1:10,000–1:15,000. Secondary antibodies used included IRDye^®^ 800CW Goat anti-Mouse IgG (LI-COR, 926-32210), IRDye^®^ 680RD Goat anti-Mouse IgG (LI-COR, 926-68070), IRDye^®^ 800CW Goat anti-Rabbit IgG (LI-COR, 926-32211), and IRDye^®^ 800CW Donkey anti-Goat IgG (LI-COR, 926-32214).

After additional washes with PBS-T and a final wash with PBS, antibody signals were visualized using the Odyssey^®^ infrared imaging system (LI-COR Biosciences).

### Copy number quantification (dPCR)

For copy number quantification, total RNA isolated from EVs or tissues was initially quantified by NanoDrop and diluted to approximately 2 ng/µL, followed by re-quantification using the QuantiFluor® RNA System (Promega). 5 ng of total RNA were reverse transcribed using a custom TaqMan™ small RNA assay (Applied Biosystems, 4398988) designed to detect the antisense strand of sh*Gapdh* (sequence: caaatccgttcacaccgacc) with the TaqMan™ MicroRNA Reverse Transcription Kit (Thermo Fisher Scientific, 10146854), according to the manufacturer’s instructions, including the optional denaturation step.

Following cDNA synthesis, EV-derived samples were diluted 500-fold, whereas tissue-derived samples were analyzed without further dilution. Digital PCR (dPCR) was performed using the QIAcuity^®^ Probe PCR Kit (Qiagen, 250101) on QIAcuity^®^ Nanoplates 8.5k (24-well; Qiagen, 250011) and run on a QIAcuity^®^ 4 Digital PCR System (Qiagen). Plates were imaged after 40 amplification cycles, and a common fluorescence threshold of 50 was applied for all analyses. Absolute copy numbers of shRNA were calculated per ng of total RNA extracted from tissue; and dilution-corrected as well as normalized to both RNA input and particle numbers for EV samples.

### Animal husbandry

Animal experiments were approved by the Swedish Animal Ethics Committee in Linköping, Sweden (permit number 14772-2023) and were conducted under the supervision of the Swedish Board of Agriculture (Jordbruksverket). All procedures were performed in accordance with national legislation and the European Union Directive 2010/63/EU governing the use of animals in scientific research. Experimental design and animal handling were carried out with the aim of minimizing animal suffering and reducing the number of animals used. Animals were euthanized using approved humane methods in accordance with institutional and national guidelines.

C57BL/6JR female mice (12 weeks of age, 18–22 g; reference SC-C57J-F) were obtained from Janvier Labs and housed at the Preclinical Laboratory (PKL), Novum, Karolinska University Hospital, Huddinge. Animals were maintained under specific pathogen-free conditions in accordance with national animal welfare legislation and the European Union Directive 2010/63/EU. Mice were acclimatized for at least 7 days prior to experimentation and group-housed in individually ventilated cages with environmental enrichment. The animals had ad libitum access to food and water and were maintained under controlled temperature (20–22 °C), humidity (45–55%), and a 12 h light/dark cycle. They were monitored daily by trained animal care staff under veterinary supervision.

### Intrastriatal injections

Intrastriatal injections were performed in C57BL/6J mice under general anesthesia with isoflurane using a stereotaxic apparatus. Bilateral injections were carried out relative to bregma at anteroposterior +1.0 mm, mediolateral ±2.0 mm, and dorsoventral −3.2 mm. EVs (1.8 × 10^10^ particles) diluted in PBS were injected in a total volume of 2 µL per hemisphere at a flow rate of 250 nL/min using a Hamilton syringe. Following injection, the needle was left in place for 5 min, retracted by 0.3 mm, and left for an additional 2 min before complete withdrawal. Burr holes were sealed with bone wax and the skin was closed with sutures. Postoperative analgesia was provided with buprenorphine (0.1 mg/kg). Mice were sacrificed 48 h after injection, and brains were harvested for subsequent dissection of the striatum.

### Statistical analysis

Statistical analyses were performed using GraphPad Prism 11. Data are presented as mean ± SD, unless otherwise stated.

For comparisons between two groups, including absolute quantification of sh*Gapdh* copy numbers per EV determined by dPCR and IC_50_ values, statistical significance was assessed using Welch’s *t*-test.

For analysis of *Gapdh* mRNA levels in mouse striatum, one-way analysis of variance (ANOVA) was applied, followed by Tukey’s multiple-comparison test to evaluate differences between groups.

For comparisons involving multiple treatment groups relative to an untreated control, one-way ANOVA followed by Dunnett’s multiple-comparison test was used.

Statistical comparisons of IC_50_ values between fusogenic conditions (VSV-G vs RVG) were performed using multiple unpaired *t*-tests with Holm–Šidák correction for multiple testing.

## Results

To prove that EVs hold potential for the delivery of shRNA, an endogenous loading platform was established. This platform relies on three crucial components: (1) an shRNA, (2) an RNA-binding protein—AGO2—fused to an EV-sorting domain (EVSD) to facilitate shRNA loading into EVs, and (3) a fusogenic protein—VSV-G—to ensure cytosolic delivery of the effector in recipient cells (**Figure 1A**). To confirm the identity and purity of isolated EVs, protein expression was analyzed by Western blot in EV preparations and their producing HEK293T cells. EVs were enriched for the canonical EV markers CD81 and ALIX, while the cellular protein β-actin was only minimally detected, indicating low levels of contamination with cellular material. Additionally, AGO2 and VSV-G were detected only in expected samples (**Supplementary Figure 1**). The contribution of each platform component to functional delivery was assessed using an shRNA targeting *GFP* (sh*GFP*). EV-mediated target knockdown in recipient cells was substantially improved when sh*GFP* loading was supported by AGO2 (65% reduction at the highest dose) and further enhanced when AGO2 contained a myristoylation site (mA) for membrane association, reaching up to 88% knockdown (**Figure 1B**). Importantly, compared to transfection of pDNA encoding sh*GFP*, EV-mediated delivery resulted in more uniform knockdown across the recipient cell population (**Figure 1C**), and without marked reductions in cell viability (**Supplementary Figure 2**). Consistent with previous reports, effective target protein knockdown occurred only when VSV-G was present on EVs (**Figure 1D**) (21, 23–26). Taken together, these observations underscore the importance of enhanced cargo loading and cytosolic release for achieving efficient EV-mediated shRNA delivery, exceeding what can be attained through passive overexpression-based loading.

**Figure 1:**
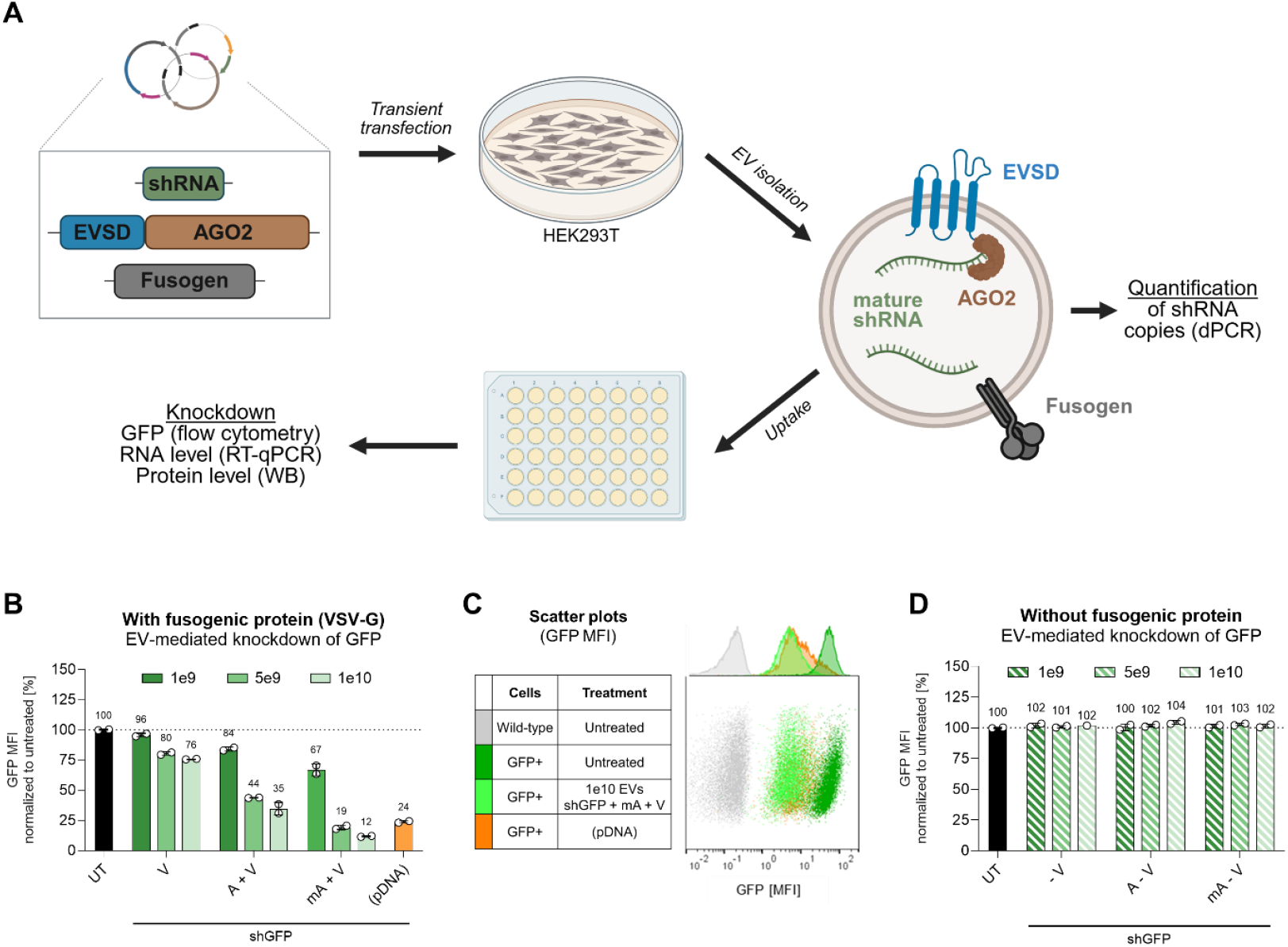
Proof-of-concept for EV-mediated shRNA delivery and functional knockdown. **A)** Schematic overview of EV production, shRNA loading, and downstream analysis. HEK293T producer cells were transfected with pDNA encoding shRNA, EVSD-AGO2, and a fusogenic protein. Engineered EVs were harvested and purified from conditioned medium, and EV-mediated target knockdown was assessed in recipient cells. **B, D)** Flow cytometry analysis of HEK293T reporter cells. Cells were treated with indicated doses of engineered EVs loaded with sh*GFP* or transfected with pDNA encoding sh*GFP*. GFP expression was measured 48 h after treatment and data are presented as MFI of the GFP-positive population, normalized to untreated (UT) cells. **C)** Representative flow cytometry scatter plots showing GFP fluorescence intensity in selected conditions from panel (B), illustrating EV-mediated reduction of GFP expression relative to control samples. EVSD, EV-sorting domain; UT, untreated; V, VSV-G; A, AGO2; mA, AGO2 with a myristoylation site; pDNA, plasmid DNA.

Moving towards more physiologically relevant conditions, we next evaluated the performance of the platform against an endogenous target (*Gapdh*) in murine neuroblastoma (N2a) cells. Potent target knockdown was achieved, with reductions of up to 87% at the protein level (**Figure 2A-B, Supplementary Figure 3**) and 96% at the mRNA level (**Figure 2C**). Notably, *Gapdh* mRNA knockdown was reproducible across independent experiments, while treatment with EVs carrying a scrambled control shRNA (shScr) had no measurable effect on *Gapdh* expression, suggesting minimal unintended effects in response to EV treatment (**Supplementary Figure 4**). To further optimize the platform, we compared the loading efficiencies of different EVSDs fused to AGO2. Specifically, the EV-sorting ability of mAGO2 (mA) was benchmarked against CD63, a widely used EV-loading scaffold, and TSPAN2, which has recently been identified as a highly efficient EV-sorting protein (27). While *Gapdh* knockdown efficiencies were comparable across these groups (**Figure 2C**), absolute quantification of sh*Gapdh* copy numbers per EV revealed pronounced differences in loading capacity. TSPAN2-AGO2 (TA)–engineered EVs carried the highest payload (3.7 copies/EV), outperforming both mA (1.03 copies/EV) and CD63-AGO2 (CA, 1.9 copies/EV) (**Figure 2D**). Based on these copy numbers, IC_50_ values were calculated for each condition (**Figure 2E**). Notably, IC_50_ values were similar across all EVSDs, including EVs in which sh*Gapdh* was loaded stochastically (VSV-G only) or facilitated by AGO2 overexpression (A + V). These results demonstrate that functional potency is primarily dictated by shRNA loading efficiency, which can be enhanced through EVSD selection and AGO2-mediated loading strategies.

**Figure 2:**
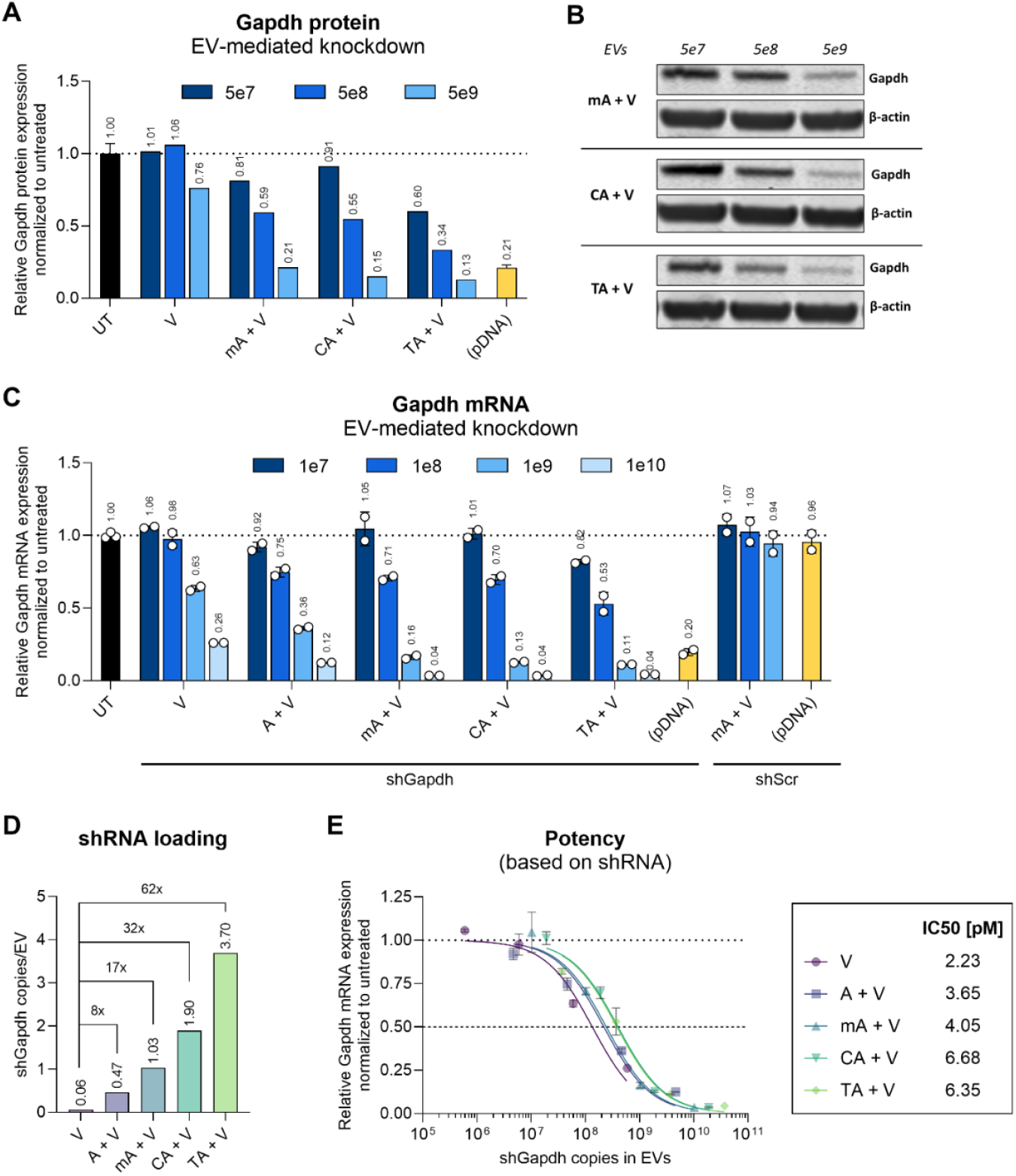
EV-mediated shRNA delivery enables efficient knockdown of an endogenous target and reveals EV-sorting–dependent loading efficiencies. **A)** Quantification of EV-mediated knockdown of endogenous Gapdh protein in N2a cells. Cells were treated with indicated doses of EVs loaded with sh*Gapdh* using different EVSDs or transfected with pDNA encoding sh*Gapdh*. Target protein knockdown was assessed 48 h after EV addition or pDNA transfection by Western blot. Gapdh signals were normalized to the β-actin loading control and are depicted relative to untreated cells. **B)** Representative Western blot images corresponding to the quantification shown in (A). Shown are selected samples from full blots provided in **Supplementary Figure 3. C)** EV-mediated knockdown of *Gapdh* mRNA in N2a cells. Cells were treated with indicated doses of EVs loaded either with shScr or sh*Gapdh* using different EVSDs or transfected with respective pDNA. Target knockdown was assessed 48 h after EV addition or pDNA transfection by RT-qPCR. Data are normalized to housekeeping gene expression and expressed relative to untreated (UT) controls. **D)** Absolute quantification of sh*Gapdh* copy numbers per EV determined by dPCR for EVs generated using different engineering strategies. **(E)** Estimated IC_50_ values derived from dose-dependent *Gapdh* mRNA knockdown (panel C) and sh*Gapdh* copy numbers per EV (panel D) for each condition. IC_50_ values were calculated by nonlinear regression using a three-parameter inhibitor-response model, with lower and upper bounds constrained to 0 and 1, respectively. EVSD, EV-sorting domain; UT, untreated; V, VSV-G; A, AGO2; mA, AGO2 with a myristoylation site; CA, CD63-AGO2; TA, TSPAN2-AGO2; pDNA, plasmid DNA; shScr, scrambled control shRNA.

Because AGO2 is an essential component of the RISC, we sought to investigate its role in EV-mediated silencing – specifically, whether its function is limited to facilitating EV loading or also contributes to target knockdown in recipient cells. To address this, AGO2 was mutated at position D597A to abolish its endonucleolytic (“slicer”) activity while retaining small RNA-binding capacity (5) (**Supplementary Figure 5**). Unexpectedly, EVs produced with TSPAN2 and the AGO2 D597A mutant (TA(D)) led to a 6.75-fold enhancement in target knockdown compared to wild-type AGO2 (**Figure 3A**). This increase in silencing potency occurred without a corresponding change in shRNA loading efficiency, as absolute shRNA copy numbers per EV were comparable between wild-type and mutant AGO2 conditions (**Figure 3B**). To further assess the contribution of AGO2 in the recipient cells, endogenous *Ago2* levels were reduced by approximately 50% using siRNA-mediated knockdown (**Supplementary Figure 6**). This partial depletion led to a pronounced decrease in EV-mediated target silencing (**Figure 3C**), indicating that endogenous Ago2 within EV-recipient cells is essential for effective target knockdown. While a contribution from vesicle-associated AGO2 cannot be fully excluded, the comparable loss of silencing activity for both wild-type and mutant AGO2 following partial Ago2 depletion argues against a direct role for exogenous, EV-associated AGO2 in target mRNA cleavage.

**Figure 3:**
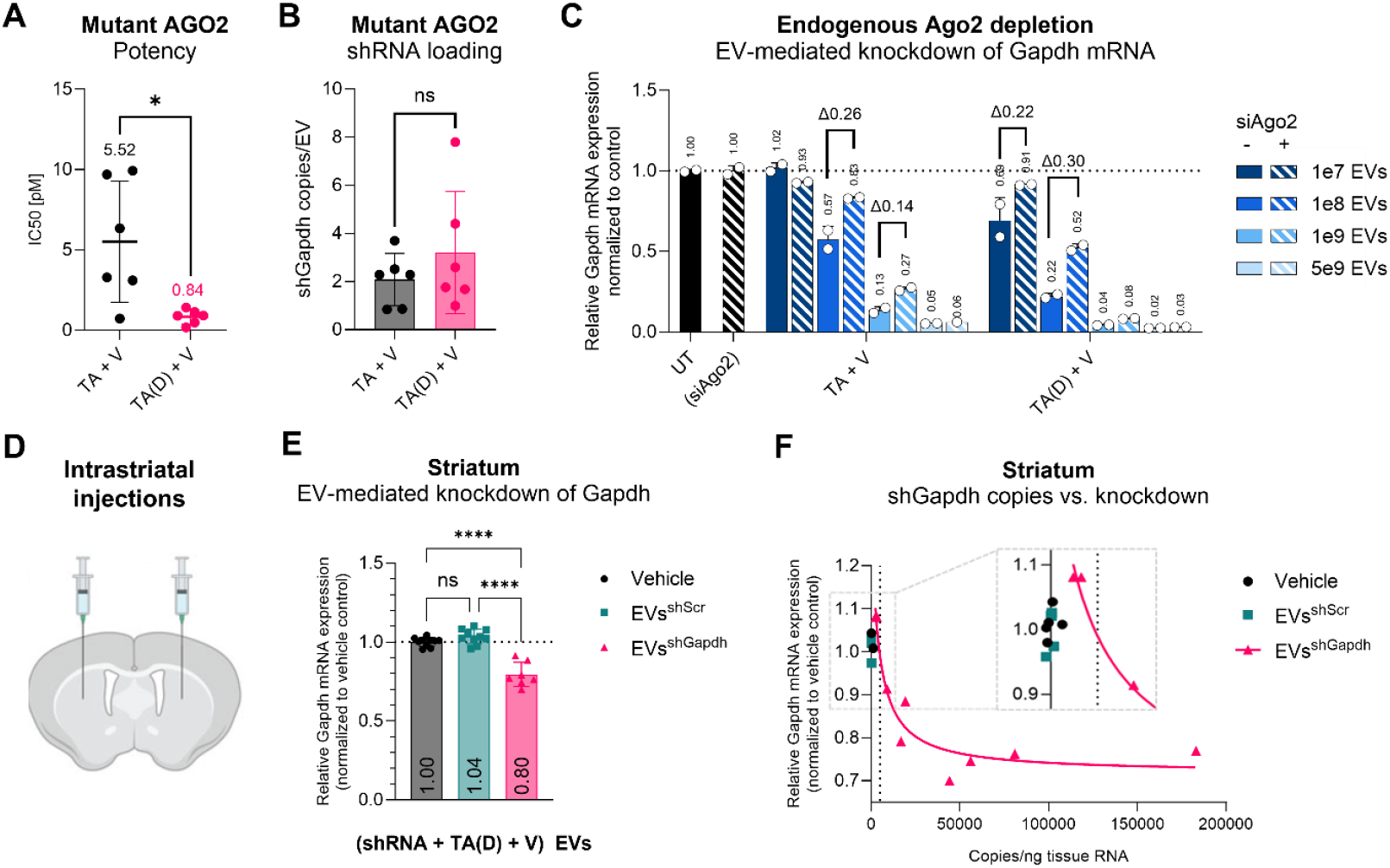
An endonuclease-deficient AGO2 mutant enhances EV-mediated shRNA silencing and demonstrates *in vivo* activity. **A)** Estimated IC_50_ values for EV-mediated *Gapdh* knockdown in N2a cells, calculated from dose-dependent target mRNA knockdown data and sh*Gapdh* copy numbers per EV for wild-type and AGO2(D597A)-mediated loading conditions. Data were analyzed by Welch’s t-test. **B)** Absolute quantification of sh*Gapdh* copy numbers per EV for wild-type and mutant AGO2(D597A)-mediated loading conditions, determined by dPCR, and analyzed by Welch’s *t*-test. **C)** Effect of partial endogenous Ago2 depletion in EV recipient N2a cells on EV-mediated *Gapdh* mRNA knockdown. Cells were reverse-transfected with si*Ago2* and treated with indicated doses of engineered EVs 24 h later. Target knockdown was assessed 48 h after EV addition, and data normalized to housekeeping gene expression and are depicted relative to untreated (UT) controls. **D)** Schematic representation of bilateral intrastriatal injections in the mouse brain. **E)** *Gapdh* mRNA levels (RT-qPCR) in mouse striatum 48 h after intrastriatal injections of EVs (1.8 × 10^10^ particles in 2 μL per injection), relative to vehicle control. Data were analyzed by one-way ANOVA with Tukey’s multiple comparison test. **F)** Correlation between sh*Gapdh* copy numbers measured in striatal tissue (by absolute quantification with dPCR and normalized to ng tissue RNA) and the extent of target knockdown shown in Figure 3E. Dashed line indicates threshold for exclusion based on detected shRNA copies. UT, untreated; V, VSV-G; TA, TSPAN2-AGO2; TA(D), TSPAN-AGO2(D597A); shScr, scrambled control shRNA; ns, not significant. P values are indicated as ^*^ P ≤ 0.05, ^**^ P ≤ 0.01, ^***^ P ≤ 0.001, ^****^ P ≤ 0.0001.

Having inadvertently enhanced the potency of the platform through the AGO2 D597A mutation, we next evaluated its efficacy *in vivo* via direct administration into the brain parenchyma. This route is particularly relevant for neurodegenerative disorders driven by region-restricted pathological gene expression, yet achieving efficient functional knockdown with RNAi therapeutics remains challenging, as limited cellular uptake, and inefficient cytosolic release necessitate high and potentially toxic doses (28). To assess EV-mediated knockdown in this context, sh*Gapdh*-loaded EVs were stereotactically injected into the striatum, and target knockdown was measured 48 h later (**Figure 3D**). EV treatment resulted in up to 30% reduction of target mRNA levels in the striatum, which was significant compared with both vehicle-treated and shScr control EVs (**Figure 3E**). To verify successful injections, shRNA copy numbers were quantified in the analyzed tissues, which allowed exclusion of two samples with low shRNA abundance and revealed a dose-dependent relationship between shRNA levels and target knockdown (**Figure 3F**). These findings demonstrate that the optimized EV-based platform supports functional gene silencing *in vivo* following direct CNS administration.

So far, EV-mediated shRNA delivery proved effective for silencing GFP in HEK293T reporter cells *in vitro*, as well as *Gapdh* in murine neuroblastoma (N2a) cells and *in vivo* in the striatum of mice. To assess the broader applicability of the platform, we next evaluated EV-mediated delivery of sh*Gapdh* across six additional recipient cell *lines in vitro*. In parallel, we examined whether functional delivery could be fine-tuned by the choice of fusogenic protein. For this purpose, VSV-G, which confers cellular tropism through binding to the low-density lipoprotein receptor (LDLR), was replaced with rabies virus glycoprotein (RVG), which binds to the nicotinic acetylcholine receptor (29) as well as the p75 neurotrophin receptor (p75_NTR_) (30). To account for differences in shRNA loading between EV preparations (**Figure 4A**), all data are reported as IC_50_ values derived from dose-response curves (**Supplementary Figure 7**). Overall, potent target knockdown was observed across all tested cell lines (**Figure 4B**). We refrained from comparing potencies between cell lines, as both experimental variables, such as cell density, and cell-intrinsic factors, including EV internalization kinetics and target gene expression, are likely to influence efficacy. Nevertheless, a general trend toward enhanced activity of RVG-pseudotyped EVs compared with VSV-G-pseudotyped EVs was observed across most recipient cell types, with the exception of C2C12 muscle cells (**Figure 4B**). These findings suggest that fusogenic protein selection may influence cellular tropism and could potentially be leveraged to tailor EV-mediated delivery.

**Figure 4:**
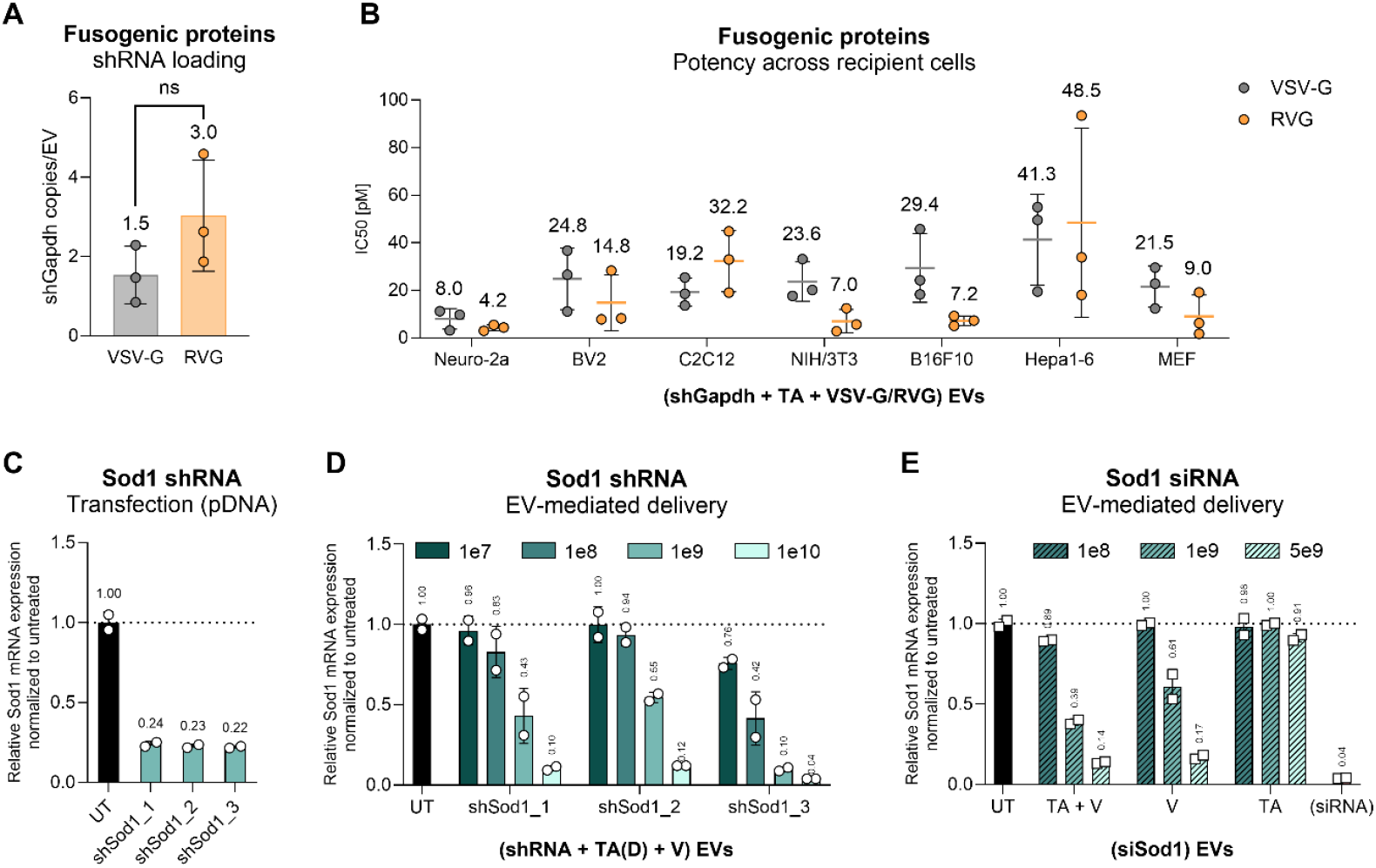
Versatile EV-mediated delivery of small RNAs across cell types, target genes, and fusogenic proteins. **A)** Absolute sh*Gapdh* copy numbers determined by dPCR for TSPAN2-AGO2 engineered EVs loaded with sh*Gapdh* and pseudotyped with either VSV-G or RVG. Data were analyzed by Welch’s *t*-test. **B)** IC_50_ values for EV-mediated *Gapdh* mRNA knockdown across different cell lines. Cells were treated with TSPAN2-AGO2 engineered EVs loaded with sh*Gapdh* and pseudotyped with either VSV-G or RVG. Target knockdown was assessed by RT-qPCR 48 h after EV addition and IC_50_ values were derived from dose-response curves (**Supplementary Figure 6**) based on copy numbers quantified in panel A. Data were analyzed using multiple unpaired *t*-tests with the Holm–Šidák for multiple-comparison correction, and no statistically significant differences were detected. **C)** RT-qPCR analysis of *Sod1* mRNA expression in N2a cells transfected with pDNA. Target knockdown was assessed 48 h after transfection relative to a housekeeping gene and normalized to untreated (UT). **D)** Quantification of EV-mediated knockdown of *Sod1* mRNA in N2a cells. Cells were treated with TSPAN2-AGO2(D597A) and VSV-G engineered EVs loaded with three distinct sh*Sod1* designs. **E**) EV-mediated delivery of si*Sod1* determined by RT-qPCR analysis of *Sod1* mRNA expression in N2a cells. Cells were treated with indicated doses of engineered EVs or transfected with si*Sod1*. Target knockdown was assessed 48 h after treatment relative to a housekeeping gene and normalized to untreated (UT). MEF, mouse embryonic fibroblasts; UT, untreated; V, VSV-G; TA, TSPAN2-AGO2; TA(D), TSPAN-AGO2(D597A); ns, not significant.

Beyond the choice of fusogenic protein, we examined the versatility of the platform for shRNAs targeting alternative mRNAs. To this end, three distinct shRNAs targeting superoxide dismutase 1 (*Sod1*) were designed. Mutations in *SOD1* are among the most prevalent genetic causes of familial amyotrophic lateral sclerosis, and the toxic gain-of-function conferred by misfolded mutant protein makes *SOD1* mRNA an attractive therapeutic target for RNAi-based intervention (31). As an initial benchmark, shRNA-expressing pDNAs were transfected into cells, resulting in comparable levels of target knockdown and thus indicating similar intrinsic potency (**Figure 4C**). In contrast, delivery of these shRNAs using the EV-based platform (TA(D) + VSV-G) revealed pronounced differences in silencing efficiency, with sh*Sod1*_3 outperforming sh*Sod1*_1 and sh*Sod1*_2 by approximately one order of magnitude (**Figure 4D**). These differences likely arise from variations in shRNA loading into EVs, suggesting that factors beyond intrinsic shRNA potency shape effective EV-mediated delivery.

Finally, we evaluated whether the platform could also support the delivery of synthetic siRNAs. To this end, a *Sod1*-targeting siRNA was transfected into EV-producing cells with or without the EV-loading component TSPAN2-AGO2 (TA) and the fusogenic protein VSV-G. Functional siRNA delivery to recipient cells was observed only when VSV-G was included and was not enhanced by TA expression (**Figure 4E**). This indicates that siRNA loading into EVs relies on a mechanism distinct from that of shRNAs and may therefore require alternative design strategies.

## Discussion

In this study, we established a versatile EV-based platform for the efficient functional delivery of shRNAs. By integrating enhanced endogenous RNA loading, fusogen-mediated cytosolic release, and adaptable cellular tropism, we demonstrated robust and reproducible gene silencing across multiple cell lines *in vitro* and *in vivo* in the mouse brain. Our findings address several bottlenecks in EV-mediated RNA delivery and provide insight into how cargo loading efficiency, endosomal escape, and RNA-interference mechanisms determine functional potency.

Comparing our findings to prior studies of EV-mediated RNA delivery remains challenging due to differences in EV source, engineering strategy, isolation and quantification workflows, and biological readouts (32). These challenges are compounded by the intrinsic heterogeneity of EVs, as well as the lack of consensus on EV dose normalization, whether based on particle number or protein content (9, 33). Moreover, a recent systematic review of EV-based small non-coding RNA therapeutics highlighted that RNA loading is rarely quantified as absolute copy numbers per EV, limiting comparability across studies and complicating translation toward standardized therapeutic dosing (34). Nonetheless, stochiometric analyses showed that individual miRNA species are present at very low abundance across EV populations, with estimates on the order of 1 copy per hundreds to thousands of vesicles (15, 35). It has been emphasized that population-averaged RNA copy numbers in EVs are typically well below one molecule per vesicle (32). This underscores the substantial improvement achieved with our loading platform, which increased the average shRNA content from approximately 0.06 copies per EV (upon shRNA overexpression alone) to 3.7 copies per EV when loading was supported by TSPAN2-AGO2 (**Figure 2D**).

Moreover, quantification of absolute shRNA copy numbers in EVs enabled calculation of functional potencies (IC_50_ values) based on the number of active molecules delivered. This analysis allowed direct assessment of EV-mediated delivery efficiency, thereby facilitating comparisons with other nanoparticle-based RNA delivery platforms, including LNPs. Although currently among the most advanced RNA delivery technologies, even optimized formulations typically display *in vitro* IC_50_ values in the nanomolar range (36, 37). By contrast, our EV-based platform achieved an average *in vitro* IC_50_ of 5.52 pM with TSPAN2-AGO2 in N2a cells, which was further reduced to 0.84 pM upon mutating AGO2 (**Figure 3A**). Direct comparisons between EVs and LNPs must nevertheless be interpreted cautiously, as these modalities differ substantially in physicochemical composition, uptake pathways and cargo stoichiometry. Despite these differences, a key mechanistic commonality between EVs and synthetic nanoparticles is endosomal entrapment, which constitutes a major intracellular bottleneck for functional delivery (38–40). Consistent with this, shRNA-mediated silencing in our system strictly required a fusogenic protein for functional delivery, underscoring cytosolic release as a critical determinant of functional EV-mediated delivery. Together, these observations suggest that although EVs currently cannot match the manufacturing simplicity and scalability of synthetic nanoparticles, their high functional efficiency per delivered RNA molecule positions them as an attractive platform for local applications (such as intraparenchymal brain injections, **Figure 3D-F**), where maximal potency and efficient cytosolic release outweigh the need for large-scale systemic dosing.

Beyond promoting endosomal release, viral fusogens provide an opportunity to modulate EV uptake and cellular tropism. Replacement of VSV-G with RVG resulted in modest, cell type dependent differences in silencing potency *in vitro*, which could indicate receptor-dependent uptake differences (**Figure 4B**). From a translational perspective, however, viral fusogens raise immunogenicity concerns that may limit repeated dosing (41). One strategy to mitigate this is the use of endogenous human fusogens, such as Syncytin-1 (42). Alternatively, targeting can be addressed through surface engineering strategies, including genetic display of antibody fragments on EV scaffold proteins (43–45), modular Fc-capture platforms enabling post-production decoration with any IgG (46), and tissue-homing peptides such as RVG29 or iRGD (47, 48). However, a key consideration across all such targeting strategies is that receptor binding and uptake do not guarantee cytosolic delivery, highlighting the importance of fusogenic mechanisms.

Enhanced silencing of target genes in recipient cells treated with shRNA-loaded EVs using endonucleolytically inactive AGO2(D597A) versus wild-type AGO2, despite comparable shRNA loading, suggests that differences in potency may arise from downstream mechanisms (**Figure 3A-B**). Notably, partial depletion of endogenous Ago2 reduced silencing for both wild-type and mutant AGO2-loaded EVs to a similar extent, indicating that endogenous Ago2 remains a key contributor to target silencing in recipient cells. Nevertheless, additional effects of vesicle-associated AGO2 cannot be excluded. In particular, AGO2(D597A) may participate in miRNA-like repression, consistent with established studies showing that slicer-deficient AGO2 supports translational inhibition and mRNA deadenylation independently of direct target cleavage (5, 49, 50). This mode of repression, however, is inherently less sequence-specific than slicer-dependent cleavage and may therefore increase the likelihood of seed-mediated off-target effects (51, 52), underscoring the need for transcriptome-wide analyses to evaluate the specificity and safety of AGO2(D597A) in EV-based RNA delivery platforms.

Our data further indicate that intrinsic RNA features influence EV-mediated delivery efficiency. Multiple shRNAs targeting *Sod1* displayed comparable silencing upon plasmid transfection, yet markedly different potencies when delivered via EVs, pointing to sequence-dependent loading effects (**Figure 4C-D**). Such behavior is consistent with growing evidence that RNA sorting into EVs is selective and regulated, governed by sequence motifs, secondary structure, and interactions with RNA-binding proteins (53–55). This observation implies that multiple factors shape effective shRNA incorporation into EVs and highlight the need for systematic investigation of sequence- and structure-dependent loading determinants. In contrast, chemically synthesized siRNAs behaved fundamentally differently. Their functional delivery depended on fusogen-mediated cytosolic release but was not enhanced by AGO2-based loading (**Figure 4E**). Unlike shRNAs, synthetic siRNAs are delivered to the producing cell as mature, fully processed duplexes that enter the cytoplasm directly without transiting through the nuclear miRNA biogenesis pathway. Consequently, they may be less likely to rely on AGO2-assisted sorting steps that facilitate incorporation of endogenously processed shRNAs into EVs. Taken together, these findings underscore that shRNA- and siRNA-based EV delivery are governed by different cellular processing routes and therefore require independent optimization strategies.

In summary, this work establishes a robust EV-based framework for shRNA delivery and provides mechanistic insights into the factors governing functional efficacy. By examining the roles of RNA loading, cytosolic release, and host RNAi involvement, our findings offer practical design principles to guide the development of EV-based RNAi therapeutics with improved potency and translational potential.

## Author Contributions

J.A.R. (Investigation, Formal analysis, Writing – original draft), G.C. (Investigation, Writing – review & editing), O.E. (Investigation, Writing – review & editing), N.K. (Investigation), D.R.M. (Investigation), X.L. (Investigation), W.Z. (Investigation, Writing – review & editing), A.M.Z. (Investigation, Writing – review & editing), H.Z. (Investigation), S.R. (Investigation), O.P.B.W. (Supervision, Writing – review & editing), I.M. (Conceptualization, Writing – review & editing), D.G. (Conceptualization, Supervision, Writing – review & editing), S.E.A. (Conceptualization, Supervision, Writing – review & editing).

## References

1. Fire, A., Xu, S., Montgomery, M.K., Kostas, S.A., Driver, S.E. and Mello, C.C. (1998) Potent and specific genetic interference by double-stranded RNA in Caenorhabditis elegans. Nature 1998 391:6669, 391, 806–811.

2. Setten, R.L., Rossi, J.J. and Han, S. ping (2019) The current state and future directions of RNAi-based therapeutics. Nat. Rev. Drug Discov., 18, 421–446.

3. Lam, J.K.W., Chow, M.Y.T., Zhang, Y. and Leung, S.W.S. (2015) siRNA Versus miRNA as Therapeutics for Gene Silencing. Mol. Ther. Nucleic Acids, 4, e252.

4. Rao, D.D., Vorhies, J.S., Senzer, N. and Nemunaitis, J. (2009) siRNA vs. shRNA: Similarities and differences. Adv. Drug Deliv. Rev., 61, 746–759.

5. Liu, J., Carmell, M.A., Rivas, F. V., Marsden, C.G., Thomson, J.M., Song, J.J., Hammond, S.M., Joshua-Tor, L. and Hannon, G.J. (2004) Argonaute2 is the catalytic engine of mammalian RNAi. Science (1979)., 305, 1437–1441.

6. Corydon, I.J., Fabian-Jessing, B.K., Jakobsen, T.S., Jørgensen, A.C., Jensen, E.G., Askou, A.L., Aagaard, L. and Corydon, T.J. (2023) 25 years of maturation: A systematic review of RNAi in the clinic. Mol. Ther. Nucleic Acids, 33, 469–482.

7. Anand, P., Zhang, Y., Patil, S. and Kaur, K. (2025) Metabolic Stability and Targeted Delivery of Oligonucleotides: Advancing RNA Therapeutics Beyond The Liver. J. Med. Chem., 68, 6870–6896.

8. Van Niel, G., D’Angelo, G. and Raposo, G. (2018) Shedding light on the cell biology of extracellular vesicles. Nature Reviews Molecular Cell Biology 2018 19:4, 19, 213–228.

9. Kalluri, R. and LeBleu, V.S. (2020) The biology, function, and biomedical applications of exosomes. Science (1979)., 367.

10. Dave, K.M., Pinky, P.P. and S Manickam, D. (2025) Molecular engineering of extracellular vesicles for drug delivery: Strategies, challenges, and perspectives. Journal of Controlled Release, 386, 114068.

11. Xu, G., Jin, J., Fu, Z., Wang, G., Lei, X., Xu, J. and Wang, J. (2025) Extracellular vesicle-based drug overview: research landscape, quality control and nonclinical evaluation strategies. Signal Transduction and Targeted Therapy 2025 10:1, 10, 255-.

12. Kamerkar, S., Lebleu, V.S., Sugimoto, H., Yang, S., Ruivo, C.F., Melo, S.A., Lee, J.J. and Kalluri, R. (2017) Exosomes facilitate therapeutic targeting of oncogenic KRAS in pancreatic cancer. Nature, 546, 498–503.

13. O’Brien, K., Breyne, K., Ughetto, S., Laurent, L.C. and Breakefield, X.O. (2020) RNA delivery by extracellular vesicles in mammalian cells and its applications. Nature Reviews Molecular Cell Biology 2020 21:10, 21, 585–606.

14. Kalluri, V.S., Smaglo, B.G., Mahadevan, K.K., Kirtley, M.L., McAndrews, K.M., Mendt, M., Yang, S., Maldonado, A.S., Sugimoto, H., Salvatierra, M.E., et al. (2025) Engineered exosomes with KrasG12D specific siRNA in pancreatic cancer: a phase I study with immunological correlates. Nature Communications 2025 16:1, 16, 8696-.

15. Chevillet, J.R., Kang, Q., Ruf, I.K., Briggs, H.A., Vojtech, L.N., Hughes, S.M., Cheng, H.H., Arroyo, J.D., Meredith, E.K., Gallichotte, E.N., et al. (2014) Quantitative and stoichiometric analysis of the microRNA content of exosomes. Proc. Natl. Acad. Sci. U. S. A., 111, 14888–14893.

16. de Jong, O.G., Murphy, D.E., Mäger, I., Willms, E., Garcia-Guerra, A., Gitz-Francois, J.J., Lefferts, J., Gupta, D., Steenbeek, S.C., van Rheenen, J., et al. (2020) A CRISPR-Cas9-based reporter system for single-cell detection of extracellular vesicle-mediated functional transfer of RNA. Nature Communications 2020 11:1, 11, 1113-.

17. Kooijmans, S.A.A., Stremersch, S., Braeckmans, K., De Smedt, S.C., Hendrix, A., Wood, M.J.A., Schiffelers, R.M., Raemdonck, K. and Vader, P. (2013) Electroporation-induced siRNA precipitation obscures the efficiency of siRNA loading into extracellular vesicles. Journal of Controlled Release, 172, 229–238.

18. Roerig, J. and Schulz-Siegmund, M. (2023) Standardization Approaches for Extracellular Vesicle Loading with Oligonucleotides and Biologics. Small, 19, 2301763.

19. Dixson, A.C., Dawson, T.R., Di Vizio, D. and Weaver, A.M. (2023) Context-specific regulation of extracellular vesicle biogenesis and cargo selection. Nature Reviews Molecular Cell Biology 2023 24:7, 24, 454–476.

20. Rädler, J., Gupta, D., Zickler, A. and Andaloussi, S. EL (2023) Exploiting the biogenesis of extracellular vesicles for bioengineering and therapeutic cargo loading. Molecular Therapy, 31, 1231–1250.

21. Liang, X., Gupta, D., Xie, J., Van Wonterghem, E., Van Hoecke, L., Hean, J., Niu, Z., Ghaeidamini, M., Wiklander, O.P.B., Zheng, W., et al. (2025) Engineering of extracellular vesicles for efficient intracellular delivery of multimodal therapeutics including genome editors. Nature Communications 2025 16:1, 16, 4028-.

22. Brown, K.M., Nair, J.K., Janas, M.M., Anglero-Rodriguez, Y.I., Dang, L.T.H., Peng, H., Theile, C.S., Castellanos-Rizaldos, E., Brown, C., Foster, D., et al. (2022) Expanding RNAi therapeutics to extrahepatic tissues with lipophilic conjugates. Nature Biotechnology 2022 40:10, 40, 1500–1508.

23. Ma, D., Xie, A., Lv, J., Min, X., Zhang, X., Zhou, Q., Gao, D., Wang, E., Gao, L., Cheng, L., et al. (2024) Engineered extracellular vesicles enable high-efficient delivery of intracellular therapeutic proteins. Protein Cell, 15, 724.

24. Zhang, X., Xu, Q., Zi, Z., Liu, Z., Wan, C., Crisman, L., Shen, J. and Liu, X. (2020) Programmable Extracellular Vesicles for Macromolecule Delivery and Genome Modifications. Dev. Cell, 55, 784.

25. Zickler, A.M., Liang, X., Gupta, D., Mamand, D.R., De Luca, M., Corso, G., Errichelli, L., Hean, J., Sen, T., Elsharkasy, O.M., et al. (2024) Novel Endogenous Engineering Platform for Robust Loading and Delivery of Functional mRNA by Extracellular Vesicles. Advanced Science, 11, 2407619.

26. Somiya, M. and Kuroda, S. (2021) Real-Time Luminescence Assay for Cytoplasmic Cargo Delivery of Extracellular Vesicles. Anal. Chem., 93, 5612–5620.

27. Zheng, W., Rädler, J., Sork, H., Niu, Z., Roudi, S., Bost, J.P., Görgens, A., Zhao, Y., Mamand, D.R., Liang, X., et al. (2023) Identification of scaffold proteins for improved endogenous engineering of extracellular vesicles. Nature Communications 2023 14:1, 14, 4734-.

28. Nikan, M., Osborn, M.F., Coles, A.H., Biscans, A., Godinho, B.M.D.C., Haraszti, R.A., Sapp, E., Echeverria, D., Difiglia, M., Aronin, N., et al. (2017) Synthesis and Evaluation of Parenchymal Retention and Efficacy of a Metabolically Stable O-Phosphocholine-N-docosahexaenoyl-l-serine siRNA Conjugate in Mouse Brain. Bioconjug. Chem., 28, 1758–1766.

29. O’Brien, B.C.V., Thao, S., Weber, L., Danielson, H.L., Boldt, A.D., Hueffer, K. and Weltzin, M.M. (2024) The human alpha7 nicotinic acetylcholine receptor is a host target for the rabies virus glycoprotein. Front. Cell. Infect. Microbiol., 14, 1394713.

30. Langevin, C., Jaaro, H., Bressanelli, S., Fainzilber, M. and Tuffereau, C. (2002) Rabies Virus Glycoprotein (RVG) Is a Trimeric Ligand for the N-terminal Cysteine-rich Domain of the Mammalian p75 Neurotrophin Receptor. Journal of Biological Chemistry, 277, 37655–37662.

31. Benatar, M., Robertson, J. and Andersen, P.M. (2025) Amyotrophic lateral sclerosis caused by SOD1 variants: from genetic discovery to disease prevention. Lancet Neurol., 24, 77–86.

32. Mateescu, B., Kowal, E.J.K., van Balkom, B.W.M., Bartel, S., Bhattacharyya, S.N., Buzás, E.I., Buck, A.H., de Candia, P., Chow, F.W.N., Das, S., et al. (2017) Obstacles and opportunities in the functional analysis of extracellular vesicle RNA - An ISEV position paper. J. Extracell. Vesicles, 6, 1286095.

33. Gupta, D., Zickler, A.M. and El Andaloussi, S. (2021) Dosing extracellular vesicles. Adv. Drug Deliv. Rev., 178, 113961.

34. Luisotti, L., Germelli, L., Piccarducci, R., Giacomelli, C., Marchetti, L. and Martini, C. (2025) Extracellular vesicles as vehicles for small non-coding RNA therapeutics: standardization challenges for clinical translation. Extracell. Vesicles Circ. Nucl. Acids, 6, 403.

35. Albanese, M., Chen, Y.F.A., Hüls, C., Gärtner, K., Tagawa, T., Mejias-Perez, E., Keppler, O.T., Göbel, C., Zeidler, R., Shein, M., et al. (2021) MicroRNAs are minor constituents of extracellular vesicles that are rarely delivered to target cells. PLoS Genet., 17, e1009951.

36. Pattipeiluhu, R., Zeng, Y., Hendrix, M.M.R.M., Voets, I.K., Kros, A. and Sharp, T.H. (2024) Liquid crystalline inverted lipid phases encapsulating siRNA enhance lipid nanoparticle mediated transfection. Nature Communications 2024 15:1, 15, 1303-.

37. Mo, Y., Keszei, A.F.A., Kothari, S., Liu, H., Pan, A., Kim, P., Bu, J., Kamanzi, A., Dai, D.L., Mazhab-Jafari, M.T., et al. (2025) Lipid-siRNA Organization Modulates the Intracellular Dynamics of Lipid Nanoparticles. J. Am. Chem. Soc., 147, 10430–10445.

38. Sahay, G., Querbes, W., Alabi, C., Eltoukhy, A., Sarkar, S., Zurenko, C., Karagiannis, E., Love, K., Chen, D., Zoncu, R., et al. (2013) Efficiency of siRNA delivery by lipid nanoparticles is limited by endocytic recycling. Nature Biotechnology 2013 31:7, 31, 653–658.

39. Mathieu, M., Martin-Jaular, L., Lavieu, G. and Théry, C. (2019) Specificities of secretion and uptake of exosomes and other extracellular vesicles for cell-to-cell communication. Nature Cell Biology 2019 21:1, 21, 9–17.

40. Chatterjee, S., Kon, E., Sharma, P. and Peer, D. (2024) Endosomal escape: A bottleneck for LNP-mediated therapeutics. Proc. Natl. Acad. Sci. U. S. A., 121, e2307800120.

41. Munis, A.M., Mattiuzzo, G., Bentley, E.M., Collins, M.K., Eyles, J.E. and Takeuchi, Y. (2019) Use of Heterologous Vesiculovirus G Proteins Circumvents the Humoral Anti-envelope Immunity in Lentivector-Based In Vivo Gene Delivery. Mol. Ther. Nucleic Acids, 17, 126–137.

42. Bui, S., Dancourt, J. and Lavieu, G. (2023) Virus-Free Method to Control and Enhance Extracellular Vesicle Cargo Loading and Delivery. ACS Appl. Bio Mater., 6, 1081.

43. Kooijmans, S.A.A., Aleza, C.G., Roffler, S.R., van Solinge, W.W., Vader, P. and Schiffelers, R.M. (2016) Display of GPI-anchored anti-EGFR nanobodies on extracellular vesicles promotes tumour cell targeting. J. Extracell. Vesicles, 5, 31053.

44. Dooley, K., McConnell, R.E., Xu, K., Lewis, N.D., Haupt, S., Youniss, M.R., Martin, S., Sia, C.L., McCoy, C., Moniz, R.J., et al. (2021) A versatile platform for generating engineered extracellular vesicles with defined therapeutic properties. Molecular Therapy, 29, 1729–1743.

45. Stranford, D.M., Simons, L.M., Berman, K.E., Cheng, L., DiBiase, B.N., Hung, M.E., Lucks, J.B., Hultquist, J.F. and Leonard, J.N. (2023) Genetically encoding multiple functionalities into extracellular vesicles for the targeted delivery of biologics to T cells. Nature Biomedical Engineering 2023 8:4, 8, 397–414.

46. Wiklander, O.P.B., Mamand, D.R., Mohammad, D.K., Zheng, W., Jawad Wiklander, R., Sych, T., Zickler, A.M., Liang, X., Sharma, H., Lavado, A., et al. (2024) Antibody-displaying extracellular vesicles for targeted cancer therapy. Nature Biomedical Engineering 2024 8:11, 8, 1453–1468.

47. Alvarez-Erviti, L., Seow, Y., Yin, H., Betts, C., Lakhal, S. and Wood, M.J.A. (2011) Delivery of siRNA to the mouse brain by systemic injection of targeted exosomes. Nature Biotechnology 2011 29:4, 29, 341–345.

48. Zhou, Y., Yuan, Y., Liu, M., Hu, X., Quan, Y. and Chen, X. (2019) Tumor-specific delivery of KRAS siRNA with iRGD-exosomes efficiently inhibits tumor growth. ExRNA 2019 1:1, 1, 28-.

49. O’Carroll, D., Mecklenbrauker, I., Das, P.P., Santana, A., Koenig, U., Enright, A.J., Miska, E.A. and Tarakhovsky, A. (2007) A Slicer-independent role for Argonaute 2 in hematopoiesis and the microRNA pathway. Genes Dev., 21, 1999.

50. Jonas, S. and Izaurralde, E. (2015) Towards a molecular understanding of microRNA-mediated gene silencing. Nat. Rev. Genet., 16, 421–433.

51. Birmingham, A., Anderson, E.M., Reynolds, A., Ilsley-Tyree, D., Leake, D., Fedorov, Y., Baskerville, S., Maksimova, E., Robinson, K., Karpilow, J., et al. (2006) 3′ UTR seed matches, but not overall identity, are associated with RNAi off-targets. Nature Methods 2006 3:3, 3, 199–204.

52. Jackson, A.L., Burchard, J., Schelter, J., Chau, B.N., Cleary, M., Lim, L. and Linsley, P.S. (2006) Widespread siRNA “off-target” transcript silencing mediated by seed region sequence complementarity. RNA, 12, 1179–1187.

53. Villarroya-Beltri, C., Gutiérrez-Vázquez, C., Sánchez-Cabo, F., Pérez-Hernández, D., Vázquez, J., Martin-Cofreces, N., Martinez-Herrera, D.J., Pascual-Montano, A., Mittelbrunn, M. and Sánchez-Madrid, F. (2013) Sumoylated hnRNPA2B1 controls the sorting of miRNAs into exosomes through binding to specific motifs. Nature Communications 2013 4:1, 4, 2980-.

54. Santangelo, L., Giurato, G., Cicchini, C., Montaldo, C., Mancone, C., Tarallo, R., Battistelli, C., Alonzi, T., Weisz, A. and Tripodi, M. (2016) The RNA-Binding Protein SYNCRIP Is a Component of the Hepatocyte Exosomal Machinery Controlling MicroRNA Sorting. Cell Rep., 17, 799–808.

55. Garcia-Martin, R., Wang, G., Brandão, B.B., Zanotto, T.M., Shah, S., Kumar Patel, S., Schilling, B. and Kahn, C.R. (2021) MicroRNA sequence codes for small extracellular vesicle release and cellular retention. Nature 2021 601:7893, 601, 446–451.

