## Supplementary Material for "Quantitative Assessment of shRNA Loading and Delivery Efficiency of Engineered Extracellular Vesicles"

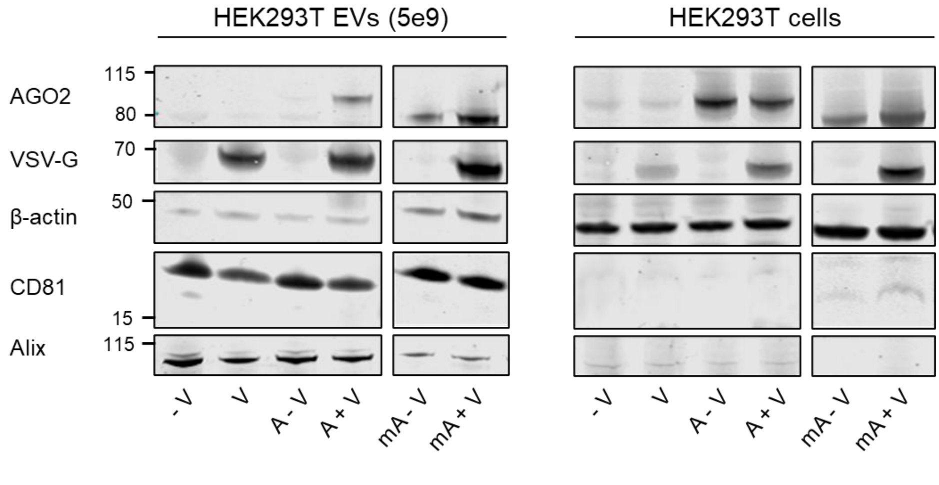


**Supplementary Figure 1: Characterization of isolated extracellular vesicles (EVs) and corresponding producer cells.** Western blot analysis of isolated EVs and their respective HEK293T producer cell lysates. EV enrichment and identity were assessed using the EV markers CD81 and ALIX. Expression of engineered components AGO2 and VSV-G was confirmed. β-actin served as a loading control for cell lysates. V, VSV-G; A, AGO2; mA, AGO2 with a myristoylation site.

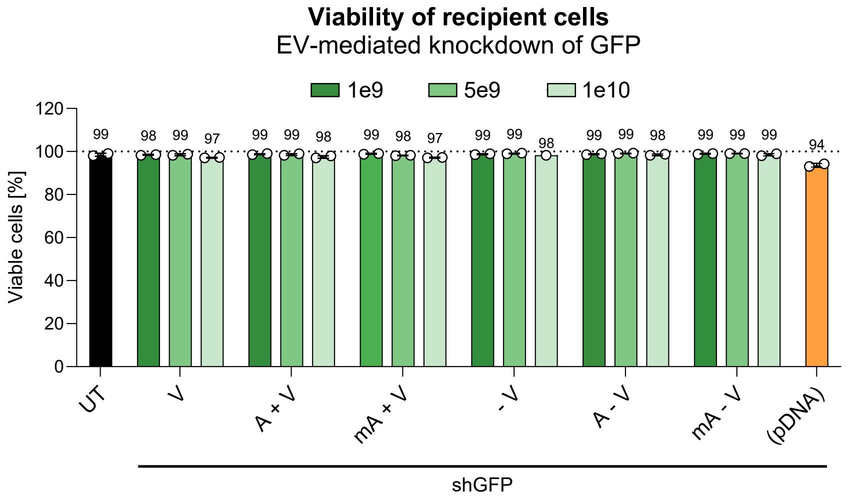


**Supplementary Figure 2: Viability of recipient cells following EV treatment and plasmid DNA transfection.** HEK293T GFP reporter cells were treated with indicated doses of engineered extracellular vesicles (EVs) or transfected with plasmid DNA (pDNA) encoding sh*GFP*. Cell viability was assessed 48 h after treatment by DAPI staining and flow cytometry. UT, untreated; V, VSV-G; A, AGO2; mA, AGO2 with a myristoylation site; pDNA, plasmid DNA.

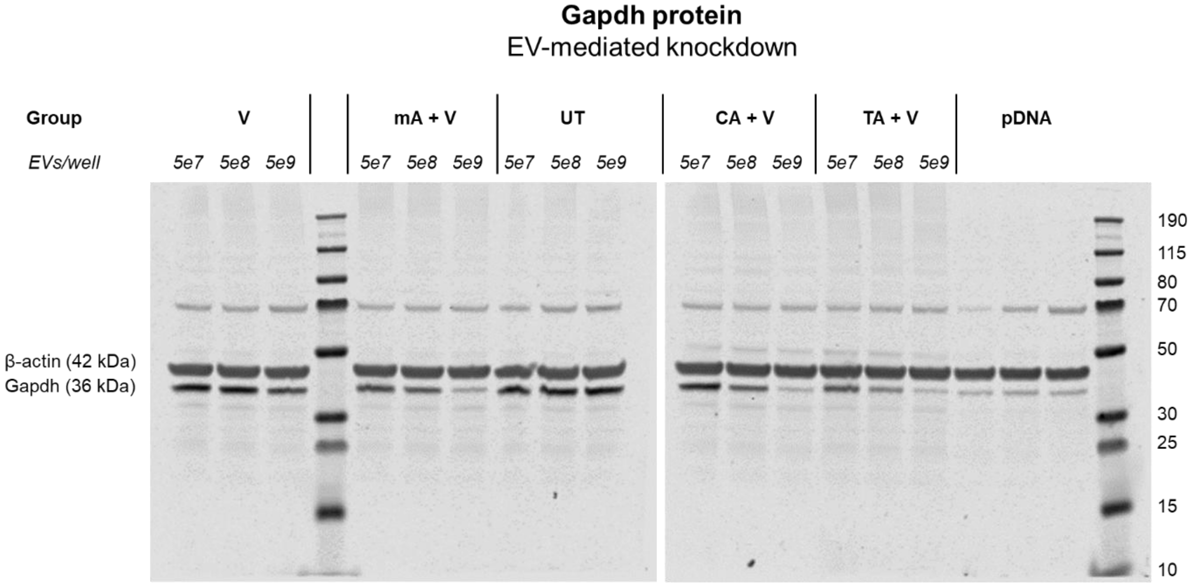


**Supplementary Figure 3: EV-mediated Gapdh protein knockdown in recipient cells.** Western blot images showing endogenous Gapdh and β-actin protein levels in murine Neuro-2a recipient cells following EV uptake in 24-well plates. Cells were treated with the indicated doses of sh*Gapdh*-loaded EVs generated using different EV-sorting domains (EVSDs) or transfected with pDNA encoding sh*Gapdh*. Cell lysates were analyzed 48 h after EV addition or pDNA transfection. β-actin served as a loading control. UT, untreated; V, VSV-G; mA, AGO2 with a myristoylation site; CA, CD63-AGO2; TA, TSPAN2-AGO2; pDNA, plasmid DNA.

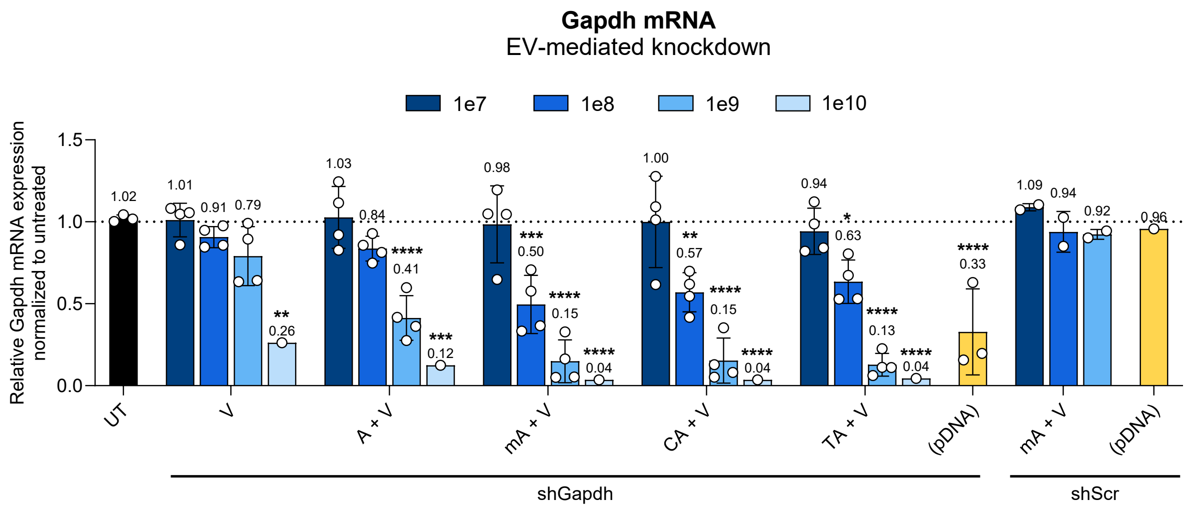


**Supplementary Figure 4: Reproducibility of EV-mediated *Gapdh* knockdown across independent experiments.** EV-mediated knockdown of *Gapdh* mRNA in murine Neuro-2a cells treated with sh*Gapdh*- or shScr-loaded EVs from independent experiments. Target knockdown was assessed 48 h after EV addition or pDNA transfection by RT-qPCR. Comparable knockdown efficiencies were observed across experiments, while EVs produced with shScr did not affect *Gapdh* expression, indicating reproducibility and specificity of EV-mediated silencing. Data represent mean ± SD from independent experiments and were analyzed by one-way ANOVA followed by Dunnett's multiple comparisons, comparing each treatment group with the untreated (UT) control. UT, untreated; V, VSV-G; A, AGO2; mA, AGO2 with a myristoylation site; CA, CD63-AGO2; TA, TSPAN2-AGO2; pDNA, plasmid DNA. P values are indicated as * P ≤ 0.05, ** P ≤ 0.01, *** P ≤ 0.001, **** P ≤ 0.0001.

### AGO2 mutation

| | | | | | | | | | | | |
|---|---|---|---|---|---|---|---|---|---|---|---|
| **Wild-type** | 1777 | TTT | CTG | GGA | GCA | GAC | GTC | ACT | CAC | CCC | 1803 |
| **D597A** | 1777 | TTT | CTG | GGA | GCA | GCC | GTC | ACT | CAC | CCC | 1803 |
| **Wild-type** | 593 | F | L | G | A | D | V | T | H | P | 601 |
| **D597A** | 593 | F | L | G | A | A | V | T | H | P | 601 |

**Supplementary Figure 5: AGO2(D597A) mutation.** Alignment showing the AGO2 coding sequence and corresponding amino-acid sequence surrounding the D597A mutation. Highlighted in red are the nucleotide substitution (1790 A→C) and the resulting amino-acid change (Aspartate to Alanine) relative to wild-type AGO2.

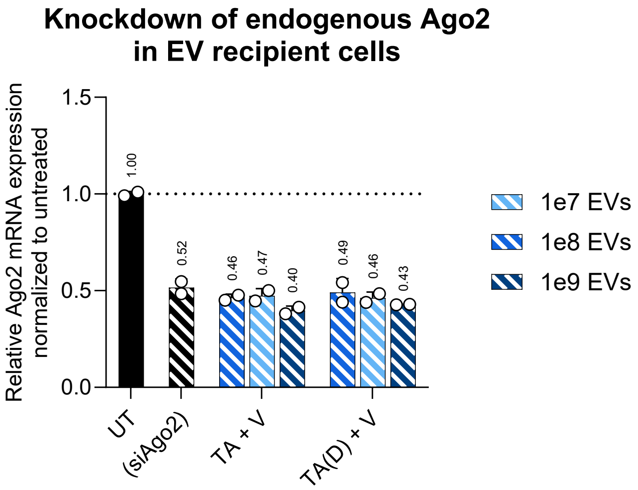


**Supplementary Figure 6: siRNA-mediated knockdown of endogenous Ago2 in EV recipient cells.** Endogenous *Ago2* levels in Neuro-2a recipient cells following siRNA-mediated knockdown and EV treatment. Cells were reverse-transfected with *Ago2*-targeting siRNA (si*Ago2*) using Lipofectamine™ RNAiMAX and subsequently treated with indicated doses of engineered EVs 24 h later. *Ago2* expression was assessed concurrently with EV-mediated target *Gapdh* knockdown (72 h post-transfection and 48 h after EV treatment) shown in Figure 3C. UT, untreated; TA, TSPAN2-AGO2; V, VSV-G; TA(D), TSPAN2-AGO2(D597A).

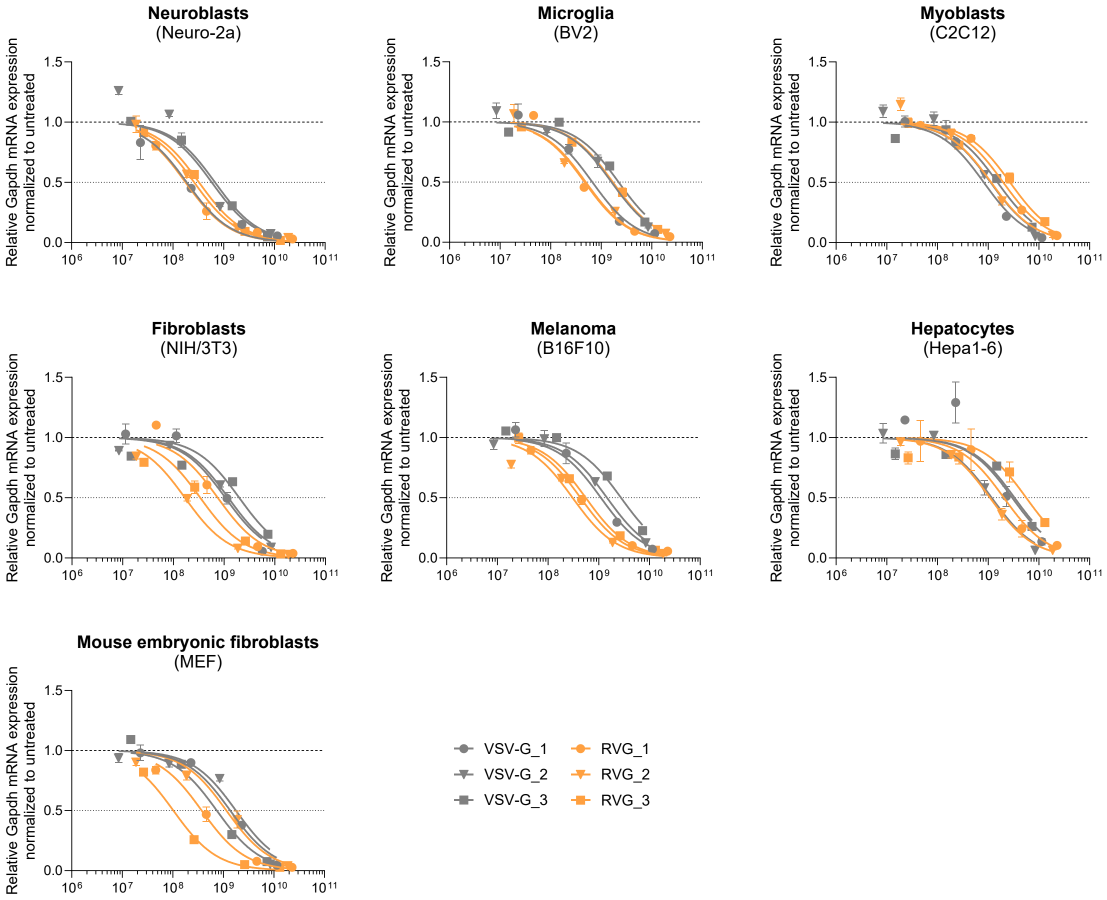


**Supplementary Figure 7: Comparison of fusogenic proteins VSV-G and RVG across multiple recipient cell types.** Dose-response curves of cells treated with sh*Gapdh*-loaded TSPAN2-AGO2–engineered EVs pseudotyped with either VSV-G or RVG. Target knockdown was assessed by RT-qPCR 48 h after EV addition. $IC_{50}$ values were derived by non-linear regression using a three-parameter inhibitor–response model with lower and upper bounds constrained to 0 and 1, respectively, and are shown in Figure 4B. Data represent three independent experiments. RVG, rabies virus glycoprotein.
